# Cryptic genes as evolutionary precursors of antimicrobial resistance: Marine bacterium *Cytobacillus oceanisediminis* encodes a sequence-divergent highly selective imipenem-degrading enzyme

**DOI:** 10.64898/2026.09.09.750327

**Authors:** Pooja Gupta, Pushplata Yadav, Jai Kishan, Saurav Gupta, Surender Shaw, Kamini Goswami, Vikash Chandra Tripathi, Kalyan Mitra, J Venkatesh Pratap, Jesu Arockiaraj, Mukesh Pasupuleti

## Abstract

The emergence of antimicrobial resistant pathogen has led to increase in mortality rate, longer hospital stays, and need for complex and expensive treatment regimens. However, it is not clear how does the bacteria develop resistance to these synthetic antimicrobial agents which have been integral part of modern medicine. One hypothesis is that non-coding cryptic genes without any vital function or positive contribution to fitness act as versatile endogenous genetic repertoires that help in bacterial adaptation and evolution of new phenotypes when conditions change. This study reports the presence of cryptic metallo beta lactamase gene in the marine bacterium *Cytobacillus oceanisediminis* isolated from the Gulf of Mannar, India. Using *in silico*, *in vitro* and *in vivo* assay, we demonstrate that the BMBL282 in *C. oceanisediminis* encodes an enzyme that can cleave only imipenem, has two characteristic domains: HXHXDH and TPGH, but does not share homology with the other reported Metallo-beta-Lactamase genes. The findings underscore the importance for comprehensive screening of cryptic antimicrobial resistance genes to fully address and mitigate the challenges posed by antimicrobial resistance. To the best of our knowledge, this is the first report of presence of cryptic gene in the Marine bacterium *C. oceanisediminis* that can degrade imipenem but not the other antibiotics of the carbapenem family.

**Importance:** One fundamental question that needs to be addressed in antimicrobial resistance evolution is, “how does bacteria get the building blocks for the genesis of antimicrobial resistant genes”?. Based on the *in silico, in vitro and in vivo* assay data, we propose that BMBL282 is a novel antimicrobial resistant cryptic gene with imipenem-degrading ability without any sequence or structure homology to reported ones and might belong to the newly proposed B3b subgroup of Metallo beta lactamase. This work clearly demonstrate that cryptic genes act as versatile endogenous genetic repertoires that help the bacterial in adaptation and evolution when required.

## Introduction

The marine environment is diverse in terms of interactions between abiotic (hydrostatic pressure, temperature, oxygen, macro-and micronutrients, and light) and biotic factors. In order to survive and adapt to complex and nutritional scarcity environment, marine microbes have a significant reservoir of phenotypically silenced junks of DNA sequences called cryptic genes (1). Cryptic genes do not express protein or RNA in the life cycle of an individual organism, even under selective pressure conditions. But the distinction is cryptic genes functional coding potential can be activated to produce phenotypic effects when it is transformed into different new species. In the natural environment, Cryptic genes could be a source of new antibiotic resistant phenotypes evolution (2).

Recent studies have indicated that Cryptic genes play a significant role in the microbial extreme capabilities to adaptation, physiology, and evolution (3, 4). Cryptic genes in the bacteria are the source or building blocks for the evolution of new phenotypes ability to produce a variety of bioactive molecules, such as enzymes, secondary metabolities, antiquorum sensing molecules etc (5). Interestingly, mathematical model suggests that there are some powerful and biologically important mechanisms which prevent the loss of these versatile endogenous genetic reservoir called cryptic genes (6).

Most bacterial species show antimicrobial resistance (AMR) through inactivation of antimicrobial agents by enzymes, especially metallo-beta lactamase (MBL) (7, 8). Bacterial strains carrying MBLs are of particular concern in healthcare settings, as they can inactivate or catalyze the hydrolysis of the “last resort” β-lactam antibiotics like carbapenems (9) which are used for treating extremely ill or harboring multidrug-resistant bacteria patients.

Recent advances in genomic sequencing techniques have led to the identification of new MBLs, and characteristics of their transfer between various pathogenic and nonpathogenic bacteria. Till date, eight families of acquired MBLs have been characterized, including imipenemase (IMPs), Verona Integron-encoded Metallo-β-lactamase (VIMs), Sao Paulo metallo-β-lactamase (SPM-1), German IMipenemase (GIM-1), Adelaide Imipenemase (AIM-)1, Seoul imipenem metallo-β-lactamase (SIM-1), New Delhi metallo-β-lactamase (NDM-1), and Dutch Imipenemase (DIM-)1 (10). IMP-type β-lactamases, which are active against imipenem, have been detected in many WHO priority pathogens, such as *Pseudomonas* spp., *Acinetobacter* spp., and members of the Enterobacteriaceae family. With the availability of a large number of bacterial genome databases, more MBL sequences from non-pathogenic, non-clinical, and obscure bacteria like marine bacteria extremophiles have been discovered in recent times (11–13).

Cryptic AMR genes add on complexity to our already silent pandemic problem of AMR. To prevent the potential therapeutic failure of next generation antibiotics, and scientifically prepared to mitigate the threat, it is utmost important that we identify and characterize cryptic AMR genes (2). If the cryptic AMR genes are not investigated, it will lead to underestimating the overall potential of antimicrobial resistance in the bacterial population and potential total antibiotic therapeutic failure in near future. Unfortunately, most of the AMR screening programs employ antimicrobial studies which are biased and recognize only resistance expressing isolates, thus totally avoiding the sensitive strains which contain the cryptic AMR (1). Given the outbreaks and the looming threat of pathogens carrying cryptic AMR genes either on chromosomes or plasmids, it is important to identify and characterize cryptic AMR genes from various sources and to better understand the mechanism and spectrum of antibiotic resistance. However, there are currently few reports on cryptic genes found in marine environments (4, 14, 15).

The present study aimed to investigate the occurrence of cryptic MBL genes in the marine bacterium *Cytobacillus oceanisediminis* isolated from the Gulf of Mannar, India. *C. oceanisediminis* cryptic MBL gene and protein phylogenetic tree analysis show that it belongs to Metallo Beta hydrolase family which included Metallo Beta lactomase, N-acyl homoserine lactone (AHL) lactonase. Furthermore, *in silico* structural analysis revealed the characteristic signature of MBL family members, such as the HXHXD and TPGH domains. Synergistic studies and disc diffusion assays with recombinant protein BMBL 282 confirmed its antagonistic effects against imipenem only with no effect on other 3^rd^ generation Beta Lactam antibiotics. Furthermore, BMBL 282 showed a better imipenem-degrading ability in the presence of Zn^2+^, which was inhibited by EDTA, as reported for other MBLs. BMBL 282 showed antagonistic effects against imipenem with FIC values of in the range of 8.01-16.08 depending upon the test bacteria. Based on the *in silico* data and biological assays, we propose that BMBL282 might belongs to a new class or i.e., B3b subgroup of MBL. Interestingly, BMBL282 showed less than 25% similarity to different B class MBL members and with no similarity to the IMP, VIM, or NDM genes. In a lung infection survival model, imipenem could not rescue mice infected with *K. pneumoniae* in the presence of BMBL282, whereas imipenem alone could rescue them. In summary, BMBL282 is a novel cryptic MBL with imipenem-degrading ability, but share no sequence or structure homology to reported MBL family members.

## Materials and Methods

### Collection of marine samples

Seawater samples were collected as previously described (16) from the Gulf of Mannar (8°28′N 79°01′E / 8.47°N 79.02°E), India in sterile tubes containing transport media (g/L NaCl 28.32; MgCl2 5.14; CaCl2 1.14; KCl 0.69; KBr 0.1; H3BO30.027; SrCl20.026; NH4Cl 0.0064; NaF 0.003 NaSiO3002; FePO40.001; yeast extract or beef extract 1.0) and packed in sterile polythene bags. The samples were transported (< 3 hours) to the laboratory on ice and processed immediately to avoid changes in the microbial composition.

### Bacterial culture conditions

The marine bacterium *Cytobacillus oceanisediminis* CDMP-24 was grown in Zobell medium at 37 °C for 24h. *E. coli* ATCC 25922 and *Klebsiella pneumoniae* ATCC 27736 was grown and maintained in MH media at 37 °C for 24h. The *Escherichia coli* DH5α strain was grown and maintained in Luria-Bertani (LB) broth at 37 °C and used as a host strain for cloning and plasmid propagation. The plasmid pET28a was used as an expression vector and *E. coli* BL21 (DE3) was used as the host cell for protein expression, when required, the medium was supplemented with 30 µg/mL of kanamycin.

### Isolation of marine bacteria from Sea samples

Marine bacteria isolation was performed as previously described by us (17). Serial dilutions of the collected samples were performed in 0.22 µm filter-sterilised seawater. The diluted samples (10^-6^) were spread on Zobell’s marine growth medium and incubated at 37 °C for two days. Colonies with different morphologies (color, appearance) were isolated and grouped. The grouped bacteria were purified to a single progeny level by streaking methods followed by passaging for 24 days on zobells agar media, and stored at -80 °C in 30 % glycerol for future use after documention.

### Bacterial identification through 16S rRNAgene amplification and sequencing

Genomic DNA was isolated from marine bacteria according to the manufacturer’s protocol (GenElute™ Bacterial genomic DNA kit mini, Sigma, Catalogue No. NA2110). The quality and quantity of the isolated DNA was analyzed using Nanodrop and agarose gel electrophoresis respectively. Amplification of the 16S rRNA gene segment was carried out with the help of a set of primers: 27F: AGAGTTTGATCMTGGCTCAG (where M can be A or C nucleotide); 63F:CAGGCCTAACACATGCAAGTC; 1387R: GGGCGGWGTGTACAAGGC (where W can be A or T nucleotide);1492R: TACGGYTACCTTGTTACGACTT, (where Y can be C or T nucleotide); 1525R:AGGAGGTGWTCCARCC, (where W can be A and T nucleotide); working concentration 1µM as reported by us elsewhere f. Briefly, a 25 µL reaction volume was set up consisting of 10 ng of marine bacteria genomic DNA, 150 ng of each primer, and 12.5 µL of Master mix (EmeraldAmp GT PCR Master Mix TaKaRa RR310). Amplification was performed in an Applied Biosystems veriti 96 well thermal cycler. The PCR cycles were pre-set to denaturation for 180 sec at 95 °C, and 30 cycles of denaturation for 60 secs at 95 °C, annealing for 60 secs at 55°C, extension for 90 secs at 72°C, then a final extension for 10 min at 72°C, 30 min at 4°C. Gel electrophoresis of the amplified 16S rRNA gene segment and elution from the agarose gel was done according to the manufacturer’s protocol (GenElute gel extraction kit, Sigma, NA1111). DNA sequencing of the eluted 16S rRNA gene segment was performed on an ABI 3730 XL sequencer. The obtained sequence was trimmed and cleaned using the DNA baser software with default settings (parameters: good bases >75%, base window= 20, QV>26, Bases with QV equal or higher than 30 were considered trusted). Chimera sequences were checked using the web-based tool DECIPHER (18) with default settings and all positions with gaps or missing data in the DNA sequence were eliminated. (19). The processed sequences were submitted to the NCBI Center for Biotechnology Information GenBank database for global records. Sequence similarity to other bacterial genomes was performed using the BLASTn program, which is available on the NCBI website. Evolutionary history was inferred using the neighbor-joining method. The percentage of replicate trees in which the associated taxa clustered together in the bootstrap test (500 replicates) is shown next to the branches. The tree was drawn to scale with branch lengths in the same units as those of the evolutionary distances used to infer the phylogenetic tree. Evolutionary distances were computed using the p-distance method (20) and expressed in terms of the number of base differences per site.

### Bioinformatic analysis of BMBL282 protein

The amino-acid sequence of BMBL282 was obtained from the UniProt database (www.uniprot.org; accession ID: A0A160M6H6). Homologous protein sequences were retrieved from the Protein Data Bank (PDB) in fasta formate. Multiple sequence alignment was performed using the Clustal Omega server (www.ebi.ac.uk). The resulting alignments were visualized and analyzed with ESPript 3.0 (https://espript.ibcp.fr) to assess sequence conservation and structural features.

The proximity of the protein (BMBL282) was analyzed by constructing a phylogenetic tree and inferred using the neighbor-joining method. The percentage of replicate trees in which the associated taxa clustered together in the bootstrap test (500 replicates) is shown next to the branches. The tree was drawn to scale, with branch lengths in the same units as those of the evolutionary distances used to infer the phylogenetic tree. The distances were computed using the p-distance method (20) and expressed in units of number of base differences per site. Evolutionary analyses were performed using MEGA11 (19). This study was conducted to determine the proximity of BMBL282 to other MBL family proteins. The analysis involved 13 MBL family protein sequences (MBL L1 type 3 *Stenotrophomonas maltophilia*, MBL AIM-1 *P. aeruginosa*, Beta-lactamase *Cronobacter sakazakii*, MBL superfamily protein *Thermus thermophilus* HB8, Lactamase B domain-containing protein *Shouchella lehensis*, N-acyl homoserine lactonase *Bacillus thuringiensis*, N-acyl homoserine lactonase AiiA *B. thuringiensis serovar kurstaki*, AIIA-like protein *Bacillus thuringiensis*, N-acyl homoserine lactone hydrolase *B. thuringiensis serovar kurstaki*, N-acyl homoserine lactonase *B. thuringiensis*), whereas G1 beta-lactamase-like protein 2 Homo sapiens was used as negative control in the analysis. All positions containing gaps and missing data were eliminated.

#### Sequence analysis

The amino-acid sequence of BMBL282 was obtained from the UniProt database (www.uniprot.org; accession ID: A0A160M6H6). Homologous protein sequences were retrieved from the Protein Data Bank (PDB) in fasta formate. Multiple sequence alignment was performed using the Clustal Omega server (www.ebi.ac.uk). The resulting alignments were visualized and analyzed with ESPript 3.0 (https://espript.ibcp.fr) to assess sequence conservation and structural features (21).

### Cloning of BMBL282 in pET28a expression vector

Genomic DNA was isolated from the marine bacteria *C. oceanisediminis* according to the manufacturer’s protocol (GenElute bacterial genomic DNA kit mini, Sigma) and PCR-based amplification of BMBL282 (846 bp) was done using *C. oceanisediminis* genomic DNA and primers BMBL282F- (5-‘CGCGGATCCTTGAAGGAAAGTTCATCT-3’) and BMBL282R (5‘-CCCAAGCTTGCCTTCCCCGTCTACATA-3’) with BamH1 and HindIII restriction sites. The primers and PCR conditions are, a 25 µL reaction volume was set up consisting of 1 ng of marine bacteria genomic DNA, 150 ng of each primer, and 12.5 µL of Master mix (Emerald Amp GT PCR Master Mix TaKaRa RR310). Amplification was performed in an Applied Biosystems veriti 96 well thermal cycler. The PCR cycles were pre-set to denaturation for 180 sec at 95 °C, and 30 cycles of denaturation for 60 secs at 95 °C, annealing for 60 secs at 55°C, extension for 90 secs at 72°C, then a final extension for 10 min at 72°C, 30 min at 4°C. The final PCR product was resolved on a 1.5 % agarose gel and Sanger gene sequencing was done using an Applied Biosystems automated sequencer. The obtained sequence was analyzed, and identification was performed using the BLAST of the National Center for Biotechnology Information.

Following agarose gel electrophoresis, the DNA fragments of interest was excised under low UV light (366 nm) to prevent DNA damage, and extracted using the GenElute™ Gel Extraction Kit (sigma Aldrich, Cat No NA1111) according to the manufacturer’s protocol. Restriction digestion of PCR products and plasmid was done in 50 µL reaction as per the manufacture instructions, 1.5 µL of BamH1 and 1.5 µL HindIII was used to digest 750 ng of PCR amplified DNA. The digested PCR products and the Plasmid DNA was placed for ligation in 10 µL reaction consisting of 1 µL ligase buffer (NEB B60025), 1 µL T4 DNA ligase (NEB M02025), 1 µL pET28a plasmid vector and 0.35 µL of the insert as per molar ratio of vector DNA, MQ added for final 10 µL volume make up. After incubation at 4°C for 12 hours, the ligation mixture was added to 200 µL of E. *coli* DH5α competent cells prepared by CaCl2 method and incubated on ice for 10 mins. A heat shock at 42 °C on dry bath for 90 seconds was given. After the heat shock, the cell was place on ice for 2 mins, followed by addition of 500 µL LB medium without antibiotic and incubate at 37°C with slow shake (100 rpm) for 45–60 min to recover the cells. After the recovery incubation of 45 mins, the cells were pelleted and spread on LB-kanamycin plates incubated for overnight at 37°C. The resultant *E. coli* pET28a-BMBL282 kanamycin resistant colonies were subjected to clone confirmation by colony PCR and restriction enzyme double digestion

### Protein Purification and confirmation

The recombinant *E. coli* BL21(DE3) carrying the BMBL282 construct was inoculated into 1 L of LB broth containing 30 μg/mL kanamycin and incubated at 25 °C under shaking conditions for 16 h at 180 RPM. Protein expression was induced with isopropylβ-D-1-thiogalactopyranoside (IPTG) concentrations (0.5 mM). The culture was harvested and resuspended in 50 mL of lysis buffer (50 mM Tris-Cl, 300 mM NaCl, and 1 tablet of protease cocktail inhibitor tablet (Roche-05892741001) and the mixture was incubated on ice for 15 min, followed by sonication in 10 sec cycles at 30 sec intervals for 25 times. The whole cell lysate was centrifuged at 10000×g for 20 min at 4 °C, and the supernatant was incubated with Ni^2+^-NTA slurry pre-equilibrated in ice-cold lysis buffer for one hour at 4 °C, loaded into the column, allowed to pass, and collected as flow-through. The Ni^2+^-NTA column was washed twice with 50 mL wash buffer 1 (300 mM NaCl, 100 mM Tris-Cl, 10 % glycerol, and 20 mM imidazole, pH 8.0) followed by 50mL wash buffer 2 (300 mM NaCl, 100 mM Tris-Cl, 10 % glycerol, and 30 mM imidazole, pH 8.0). The proteins were eluted with an elution buffer (300 mM NaCl, 100 mM Tris-Cl, 10 % glycerol, and 300 mM imidazole, pH 8.0). To remove excess salts and imidazole, the protein was dialyzed with a 10 kDa dialysis membrane (Merck, NJ, USA) in dialysis buffer A (50 mM Tris-Cl, 300 mM NaCl, and 10 % glycerol) for 3 h and replenished with dialysis Buffer B (50 mM Tris-Cl, 100 mM NaCl, and 5 % glycerol) for 4 h at 4 °C. After dialysis, the protein was concentrated using 10kDa cut off Amicon protein concentrator and concentrated protein was aliquoted and stored at – 80 °C for further studies. Protein samples were separated using sodium dodecyl sulfate–polyacrylamide gel electrophoresis (SDS–PAGE) and transferred onto nitrocellulose membranes using a Bio-Rad transfer system (Bio-Rad, USA). The membranes were blocked in 5 % nonfat milk for 1 h at room temperature and then incubated with a primary anti-His antibody (1:1000; PA5-51700, Invitrogen, USA) at 4 °C overnight. After incubation, the membranes were washed three times with Tris-buffered saline-Tween 20 (10 mM Tris, 150 mM NaCl, and 0.05 % Tween 20) and further incubated with a secondary anti-rabbit IgG antibody (1:5000, Cell Signaling Technology, USA) for 1 h at room temperature. Protein bands were stained by incubation with enhanced chemiluminescence (ECL) solution (Millipore, USA) and visualized using the ChemiDoc System (Bio-Rad).

### Agar disc-diffusion method

Test antibiotic loaded discs (approximately 6 mm in diameter, Himedia, India) were soaked in 15 µl of BMBL282 (250 µg/ml) in 50 mM MOPS buffer (pH 7.5) for 12 hours at 37 °C on gentle shaking. LB agar plates were spread with 1-2×10^6^ (0.5 MacFarland) inoculum of the test bacteria and BMBL282 treated antibiotic discs were placed on the agar plates. Disk-diffusion susceptibility tests were performed with the following antibiotics Imipenem, Meropenem, Doripenem, Ertapenem, and Penicillin-G (10 μg/disc) of standardized concentration were placed on the agar surface. The petri dishes were incubated for 24 h at 37 °C and zones the inhibition diameters were measured and reference to the control i.e untreated disc of same antibiotic.

### Determination of fractional inhibitory concentration (FIC) and FIC Index (FICI)

The checkerboard broth microdilution method was used to determine the synergy between the BMBL 282 and antibiotics (Imipenem or Meropenem), as described previously (22). The starting highest concentrations of the antibiotics were 25µg/ml and 250 µg/ml for BMBL282. A two-fold serial dilution of the antibiotic and two-fold serial dilutions of BMBL 282 was prepared for each combination tested. After 12 hours of incubation of antibiotics with BMBL282, bacterial inoculum (1-2 × 10^5^ CFU/ml) was added and incubated for further 18 hours at 37 °C. After incubating, 15 μl of filtered sterilized 0.05 % resazurin solution was added to each well and further incubated for 1 h. MIC was defined as the lowest concentration showing no color change, which exhibited complete inhibition of growth.

The checkerboard method is used to calculate the fractional inhibitory concentration (FIC) index (FICI). The FIC was derived from the lowest concentration of the antibiotic and imipenemase (BMBL 282) combination, showing no color change of resazurin. Interactions were classified as synergistic (ΣFIC ≤ 0.5), additive (≥ 0.5–1.0), indifferent (≥1.0–≤4.0), or antagonistic (ΣFIC > 4.0).

The FIC value for each agent was calculated using the following formula: FICI = FIC (antibiotic) + FIC (BMBL 282), where FIC (antibiotic) = MIC of antibiotic in combination/MIC of antibiotic alone. FIC (BMBL 282) = MIC of BMBL 282 in combination/MIC of BMBL 282.

### β-Lactamase Activity Assay

Imipenem cleavage was measured as described previously elsewhere (23). Briefly, the β-lactamase activity of purified BMBL 282 was determined by monitoring the decrease in β-lactam absorbance resulting from hydrolytic cleavage at every one hours for 6 hours. The reaction was performed at 25 °C in a mixture containing purified BMBL 282 (500 µg/mL), 50mM MOPS buffer (pH 7.5), 6.814 µg/mL ZnCl₂, and imipenem (100 µg/mL). Imipenem degradation was also assessed in the presence of 37 µg/mL EDTA to evaluate the effect of metal chelation. The decline in absorbance associated with imipenem hydrolysis was recorded spectrophotometric ally at 300 nm on Tecan infinite 200 pro plate reader, USA.

Steady-state kinetic parameters of purified BMBL282 were determined spectrophotometrically by monitoring imipenem hydrolysis. Reactions were carried out at 25 °C in 50 mM MOPS buffer (pH 7.5), 500 μg/mL BMBL282 supplemented with 6.814 µg/mL ZnCl₂, using imipenem concentrations ranging from 0.05 to 0.8 mM. Hydrolysis of the β-lactam ring was measured as a decrease in absorbance at 300 nm (23). Kinetic parameters were determined by Michaelis-Menten analysis using GraphPad Prism5.

### Agar diffusion assay with Agrobacterium tumefaciens A136 and Chromobacterium violaceum 026

To determine the quorum sensing inhibition ability of BMBL 282, a range of AHL molecules, including short- and long-chain AHL, were tested using *Agrobacterium tumefaciens* A136 and *Chromobacterium violaceum* 026 as test organisms as reported elsewhere with slight modification (24). For the bioassay, 20 µl of reaction mixture consisting of 0.1 µM of BMBL282 and 0.2 mM various AHL substrate in 50 mM Tris-Cl (pH 7.4) was incubated at 37°C for 12 hrs with gentle agitation in the presence and absence of 100 µM ZnCl2. Briefly, the biosensor strains were grown overnight at 30°C in LB broth supplemented with Spectinomycin (50 μg/mL) and Tetracycline (4.5 μg/ml) for *A*. *tumefaciens* A136 and Kanamycin *(25* μg/ml) for *C. violaceum* 026. Fresh LB agar plates were prepared by inoculating the overnight grown culture of *A. tumefaciens* A136 supplemented with X-gal (250 μg/ml) or *C. violaceum* 026 and wells (5mm) were created with the help of *Accu Sharp* Biopsy punch followed by loading of 15 µl reaction mixture. The plates were further incubated for 24 hrs at 30°C. AHL dissolved in 50 mM Tris-Cl (pH 7.4) were used as positive and 50 mM Tris-Cl (pH 7.4) as negative control.

### Normalized β-Galactosidase activity with Agrobacterium *tumefaciens* A136

The β-galactosidase activity assay was performed as previously reported with some modifications (25). BMBL 282 (1µg/ml) was added to 175µl LB broth supplemented with AHLs C6 to C14 HSL (0.2 mM) with their oxo group and PIPES buffer (100mM pH 7.4) and incubated for 24 h at 37 °C (180 RPM). After incubation, 10 µL of the reaction mixture was added to 100 µL of 1:100 diluted overnight grown culture of *Agrobacterium tumefaciens* A136 in LB broth supplemented with X-gal (250 µg/ml), followed by incubation at 30 °C for 12-16 hrs. The cells were pelleted by centrifugation, resuspended in 200 µL DMSO, vortexed for 1 min at high speed, and centrifuged at 8000 RPM for 3 min. The supernatant was collected, and the absorbance was measured at wavelengths of 492 nm and 630 nm. β-galactosidase activity was calculated using the following formula:

Normalized β-galactosidase activity= (0.653×OD492-OD630)/(0.267 ×OD630-OD492)

### Protein and ligand preparation for docking studies

A three-dimensional (3D) structural model of *Cytobacillus oceanisediminis* BMBL282 was generated using the Chai Discovery web server (https://lab.chaidiscovery.com/) (26). The predicted active site of the modeled protein was further validated using the web-based tool HotSpot Wizard 3.0 (27). Two zinc ions were incorporated by superposing the BMBL282 model with NDM1 crystal structure (PDB ID: 5YPL; 20.5% sequence identity; later used as control in docking process) with RMSD 2.2 in the PyMOL (28, 29). The finalized protein structure, obtained in PDB format, was converted to PDBQT format using MGLTools version 1.5.7 (30) for molecular docking studies. Annotated antibiotics, imipenem (PubChem ID: 104838) were retrieved from the PubChem database. The ligand structures were downloaded in SDF format and converted to PDB format using PyMOL (29). Prior to docking, polar hydrogen atoms were added and Gasteiger charges were assigned using AutoDock tools. The prepared ligands were then saved in PDBQT format for subsequent docking analyses (30).

#### Molecular docking and analysis

Molecular docking of the BMBL282 protein with the selected ligand compounds was carried out using AutoDock version 1.5.7 (30). Prior to docking, a grid box was defined around the predicted active site to ensure appropriate sampling of the binding region. The protein structure was prepared by assigning Kollman charges and adding polar hydrogen atoms using AutoDock Tools, after which the structure was saved in PDBQT format (31). A three-dimensional grid box with dimensions of 50 × 50 × 50 Å were generated and centered on the active site of the BMBL282 protein. Grid and docking parameter files (GPF and DPF) were created using AutoDock Tools. Docking were performed using the Lamarckian Genetic Algorithm, which generated multiple binding conformations for each protein-ligand complex (30). For each docking experiment, ten independent conformations were generated to evaluate binding energies. The protein–ligand complex exhibiting the lowest binding energy and matching conformation with control, was selected for further analysis. The resulting docked complexes were visualized using Protein Plus server (https://proteins.plus) (32).

The use of molecular docking studies allows for a thorough investigation of the interactions between ligands and proteins. Molecular docking tool AutoDock 4.2 was used to find the better binding affinity and interactions between established carbapenem antibiotics against the BMBL282 protein, and approximately ten different dockings were conformed with the receptor. The chosen ligand was docked from MGLTools version 1.5.7 (30), superposed with hydrolyzed imipenem bond NDM structure with RMSD 2.2 Å (5YPL) and calculated binding energy from PRODIGY server (https://wenmr.science.uu.nl/prodigy/lig)(33). The molecular binding interaction of ligands with complex protein molecules is visualized from Protein Plus server (https://proteins.plus/)(32).

### Stability of BMBL282 in Lung Lavage Fluid

Balb/c mice (15-25g) used for this study were obtained from the National Laboratory Animal Ethics Committee (IAEC/2023/45/Renew-2/Dated-30/06/2025). Mice were anesthetized using a ketamine/xylazine cocktail solution (100 mg/ml and 20 mg/ml, respectively). Following anaesthesia, the thoracic cavity was surgically opened to expose the trachea, which was then carefully cannulated. The lungs were filled with 700μl sterile phosphate-buffered saline (PBS; pH 7.4), and the bronchoalveolar lavage fluid was collected using 1 mL syringe as reported elsewhere (34). The collected fluid was centrifuged at 1,500 RPM for 10 minutes at 4 °C to remove cellular debris, and the clarified supernatant was used immediately for further analysis. Purified His-tagged BMBL282 (1mg/ml) was incubated with lung lavage fluid (1:1, v/v) at 37 °C for 6 hrs at shaking condition. Control samples included BMBL282 alone and lung lavage fluid alone incubated under identical conditions. Reactions were terminated by addition of Laemmli sample buffer with β-mercaptoethanol and heating at 95 °C for 5 min. Samples were immediately resolved on 12% SDS–PAGE to avoid any sample damage and later transferred onto a PVDF membrane. The membrane was probed with anti-His tagged primary antibody (Abclonal, AE003)) followed by anti-mouse HRP-conjugated secondary antibody (CST,7076S) and signals were detected using enhanced chemiluminescence. Protein stability was assessed by comparing band intensity and integrity between treated and control samples.

### In vivo lungs experiment

The imipenan degrading activity of BMBL282 in complex environments was evaluated using *K.pneumoniae* a lung infection model. The Balb/c mice (15–25 g) used in this study were obtained from the National Laboratory Animal Ethics Committee (IAEC/2023/45/Renew-2/Dated-30/06/2025). Before initiating the experiment, the mice were acclimatized to laboratory settings for 3 days. Subsequently, the mice were housed in IV cages in an animal house with appropriate temperature and food. The mice were made neutropenic by administering cyclophosphamide via the IP route at 96 and 24 hours before bacterial infection at 150 and 100 mg/kg body weight, respectively. Just 24 hours before bacterial infection *K.pneumoniae* ATCC 27736 strains were grown to mid log phase and adjusted to an optical density at 620 nm (OD620) of 0.4. Bacteria were then harvested by centrifugation (10000 RPM for 10 min at 25 °C) and resuspended in sterile 1X PBS at a final concentration of 1-2 × 10^7^ CFU/ml, based on optical density. On the day of bacterial infection, the mice were divided (10 per group) and anesthetized using a ketamine/xylazine cocktail solution (100 mg/ml and 20 mg/ml, respectively). Once the mice were properly anesthetized, they were placed in the supine position with the help of their incisors on a slanted platform. Using blunt-end forceps, the tongue of the mice was carefully pulled out and 100 µl of bacterial solution (1-2 × 10^7^ CFU/ml) was introduced into the trachea of the mice using the Microsprayer Aerosolozer (PennCentury, Inc.; Model IA-1C and FMJ-250 high-pressure syringes). The aerosolizer was not inserted beyond the kink/bend region but was aligned with the incisors. The aerosolizer was slowly removed and the mice were carefully transferred to their respective cages for recovery. As a non-infected control, intratracheal instillation was performed using normal saline in the same manner. To assess the efficacy of BMBL282, mice were administered a dose of 250 µg/kg and imipenem 5 mg/kg intratracheally at 12 and 36 h post-infection. The survival of the mice was calculated for 7 days to determine the efficacy of the enzyme.

### Statistical Analysis

The experimental results were analyzed by calculating the mean ± standard deviation (SD) of triplicate experiments. Statistical significance was determined using one-way ANOVA and Tukey’s test in sigma plot.

## Results

### Collection and Isolation of marine bacteria

Marine bacteria isolation and selection were performed as previously described (35) from the marine water samples collected from Gulf of Mannar, India (Supplementary Fig. S1A). In the first week, more than 50 colonies per plate from the 10^-6^ dilution appeared on the plate, however only 30 different isolates survived the 24 days of continuous culture under laboratory conditions. Cultures that were able to grow on Zobell’s agar even after 24 days were selected and glycerol stocks were prepared (Fig. 1A) as they were sustainable for culture in the laboratory condition. As reported by other, most of the isolated colonies did not survive the long period of purification in our cases also. Continuous culturing of marine bacteria isolate in artificial media is very difficult, even today, as the specific requirements for marine bacterial species are not well understood (36, 37).

**Fig 1:**
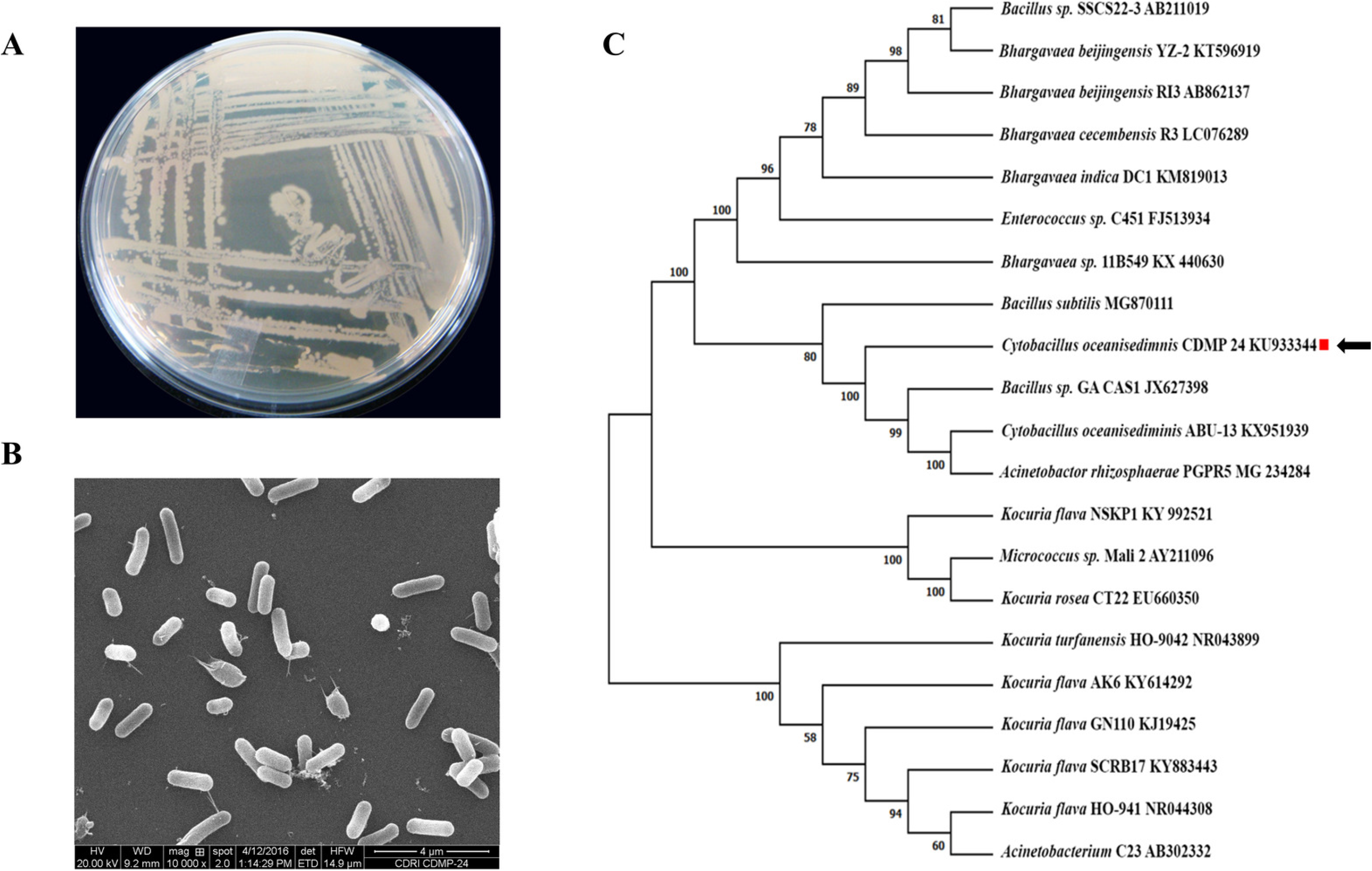
*Cytobacillus oceanisediminis* (CDMP24) culture and identification (A) Colony growth on Zobell agar media (B) Electron microscopy image (C) N-J method of Phylogenetic analysis of *C. oceanisediminis* using 16’srRNA with bootstrap of 1000.

It is well known that marine bacteria can grow at moderate to high salt concentrations. To rule out cross-contamination or mismatch, all the isolates were subjected to the salt tolerance assay, as terrestrial bacteria do not survive in medium with high sodium concentration. The average salinity of the Gulf of Mannar, Bay of Bengal, India, water is

2.5 % to 3.5 %, and this salinity varies according to the depth level (38). To determine the salt tolerance level of all marine isolates, we cultured the isolated bacteria at varying sodium chloride concentration ranging from 0-20 % with an increment of 2 % (w/v). Among all the isolates, one isolate, designated as CDMP 24, survived up to 10 % salt concentration and was selected for further study (Supplementary Fig. S1*B*).

### Marine bacterial isolate characterization and identification

To avoid redundancy and repetition of work, we focused on the identification and characterization of the marine bacterial isolate CDMP 24 using both classical and molecular characterization methods. Initially, Gram staining and scanning electron microscopy (Fig. 1B), followed by PCR amplification and sequencing of the 16S rRNA gene using universal primer sets (27F and 1492R; 519F and 1389R) (SI Appendix table 1) was performed. In comparison to terrestrial bacteria, marine bacteria species genomes are not well characterized. Hence to rule out false identification, two sets of universal primers were used and amplicon obtained after PCR amplification with both the primers sets was found to have a size of around 1.5 kb (Supplementary Fig. S1*C*). The PCR amplicons were subjected to Sanger sequencing to obtain the exact DNA sequence. Before the BLAST analysis, the obtained sequence was checked for contigs and inconsistencies using the web-based tools DECIPHER and DNA Baser (18, 39). Interestingly, when blastp analysis was done with the both sequences separately, the analysis results obtained in both cases were similar and indicated that bacteria CDMP24 isolate is *Cytobacillus oceanisediminis.* BLAST and phylogenetic tree analysis showed that the isolate is *C. oceanisediminis* which shares a close evolutionary relationship with bacteria belonging to the genus *Bacillus* but was readily distinguishable from its closest phylogenetic neighbors (Fig.1C). The obtained 16S rRNA sequence of *C.oceanisediminis* was submitted to NCBI (Accession number KU933344.1) (Supplementary Fig. S1D).

*C.oceanisediminis* is a Gram-positive, spore-forming, rod-shaped aerobic bacterium that belongs to the recently proposed novel genus *Cytobacillus* (Fig. 1C) (40). It was intially isolated from a marine sediment sample from the South Sea in China and has been shown to have metal resistance and bioremediation activity (41) Nabil-Adam later reported the existence of *C.oceanisediminis* from Abu-Qir Bay in Alexandria, Egypt and shown that ethanolic extract contains three different compounds (3-hydroxytyrosol, benzoic acids, and Rosmarinus) which showed hepatoprotective activity (42). Our data are in agreement with the study results of Zhang *et al.*, who showed that *C.oceanisediminis* grows well under variable conditions ranging from 4–45 °C (optimum 37 °C), pH 6–10 (optimum pH 7.4), and 0–13 % (w/v) of NaCl concentrations (41).

### Bioinformatics analysis of BMBL282 protein

To identify MBL cryptic genes, PCR amplification using different MBL primer sets reported in the literature (SI Appendix table 1) was done. Out of them one primer showed amplification of around 848bp. Later DNA sequence obtained by us using sanger sequence method was translated to protein sequence using online software (https://web.expasy.org/translate/). The translated amino acid sequence was 282 amino acids long (here after referred as BMBL282, B for the bacillus species, MBL for metallo-β-lactamase, 282 number of amino acids), showed to contain highly conserved zinc-binding HXHXDH signature domain of the metallo-β-lactamase (MBL) fold superfamily and TPGH domain (Fig. 2*A* and *B*). Further, In silico location locus identification studies revealed that BMBL282 was located on the chromosome complementary strand (base pair 410447 to 411295; locus tag=A361_02135) and was annotated as MBL-fold metallohydrolase (protein_id=AND37992.1) in *C.oceanisediminis* 2691, (complete sequence NCBI GenBank accession CP015506.1) (Supplementary Fig. S2A).

**Fig 2:**
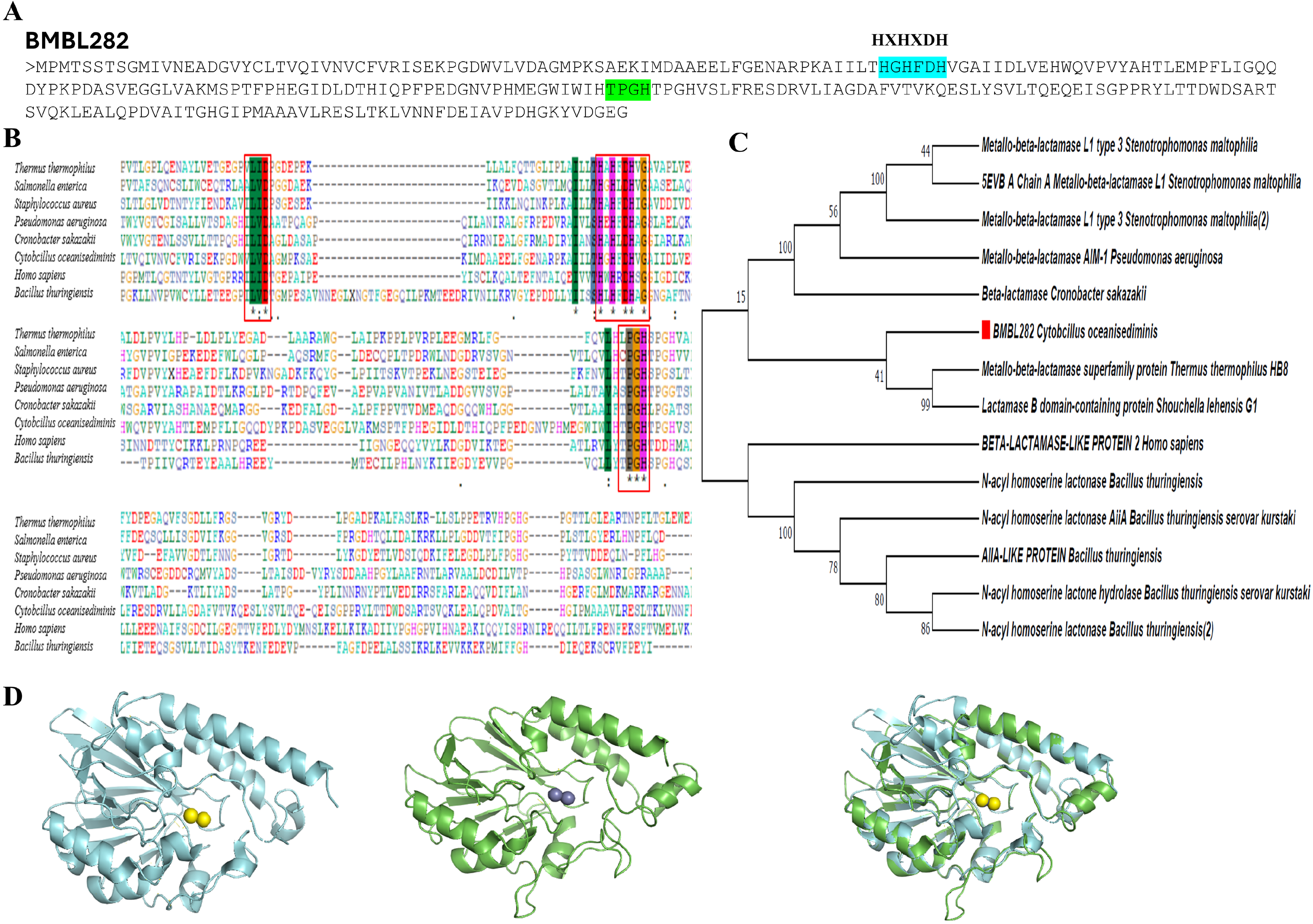
Molecular and structural analysis of BMBL282 (A) BMBL282 protein sequence (B) ClustalW alignment of BMBL282 showing the conserved HXHXDH and TPGH domain. (C) N-J method of Phylogenetic analysis of BMBL282 showing clustering with metallo-beta-hydrolases. (D) Homology Model of BMBL282 (left blue) with Metallo-beta-Lactamase Adelaide IMipenmase 1 (4AX0) from *Pseudomonas aeruginosa* (middle green) and superimpose (right).

To get an indication about potential substrate or group to further explore, phylogenetic analysis was performed using the BMBL 282 amino acid sequences. The results showed that BMBL 282 clustered in the metallo-beta hydrolase group, with a close evolutionary relationship to AHL degrading enzymes lactonase and MBL. Interestingly, our protein multiple alignment results revealed that BMBL282, even though clustered with the MBL class, it did not group with the other reported MBLs like NDM, VIM, and IMP clusters (Fig. 2*C*).

For better understanding, we tried to generate the crystal structure of BMBL282, but multiple attempts to crystallization failed with no useful data being generated (Supplementary Fig. S4). Given this *in-silico* approaches were employed to continue protein-ligand interaction studies. Initial structural modeling was carried out using an AlphaFold-predicted model; however, this model adopted a closed conformation that was unsuitable for ligand binding. To further understand, a 3D structural model of BMBL 282 was generated by submitting its amino acid sequence to the SWISSMODEL online server (https://swissmodel.expasy.org/), an automated software that calculates models based on known structural templates and sequence structure alignments. The solved structure of Metallo-beta-Lactamase Adelaide IMipenmase 1 (4AX0) from *Pseudomonas aeruginosa* was selected as the template based upon the RMSD (0.493) value and top hit. Next, the structural model was aligned to the structure of imipenemase using PyMOL Molecular Graphics System (Fig. 2D).

Despite the MBL sharing less than 25 % homology among them, the active site architecture is quite superimposable with one another, has unique alpha-beta-beta alpha fold and a flexible loop to accommodate wide range of substrates (43). The structural overlap indicates that the two enzymes i.e., BMBL 282 and *P.aeruginosa* imipenemase, superpose well and share similar motifs and active sites, zinc binding, catalytic sites, and other coordinating structure residues. In short, *in silico* 3D structural model suggested that *C.oceanisediminis* BMBL 282 might have the metallo beta-lactamase activity.

### BMBL282 Cloning expression and protein purification

As BMBL 282 did not group very well with the other known MBLs, it was interesting to explore the potential function of this gene BMBL 282. To achieve this objective, an 848bp amplicon encoding BMBL 282 was ligated into the pET28a vector digested with BamHI and HindIII restriction enzymes, with a His tag (six residues) at the C-terminus. The pET28a -BMBL282 construct was transformed into *E. coli* BL21, and recombinant expression of BMBL 282 was induced with 0.5mM IPTG at 25°C for 16 h. The expressed soluble protein was purified using Ni-NTA column and eluted with 300 mM imidazole solution. The size of the purified N-terminal 6×His-tagged recombinant protein was estimated to be approximately 33 kDa by SDS-PAGE, which corresponds to a predicted molecular weight of 32 kDa (http://www.expasy.org.tools/) (Fig. 3). The difference between the observed and expected molecular weights was attributed to the 6×His tag. The purified recombinant protein was stored at -80 °C for further downstream processing. To confirm the recombinant nature, western blot analysis was performed using an anti-His-tag antibody (Supplementary Fig. S2*B*).

**Fig 3:**
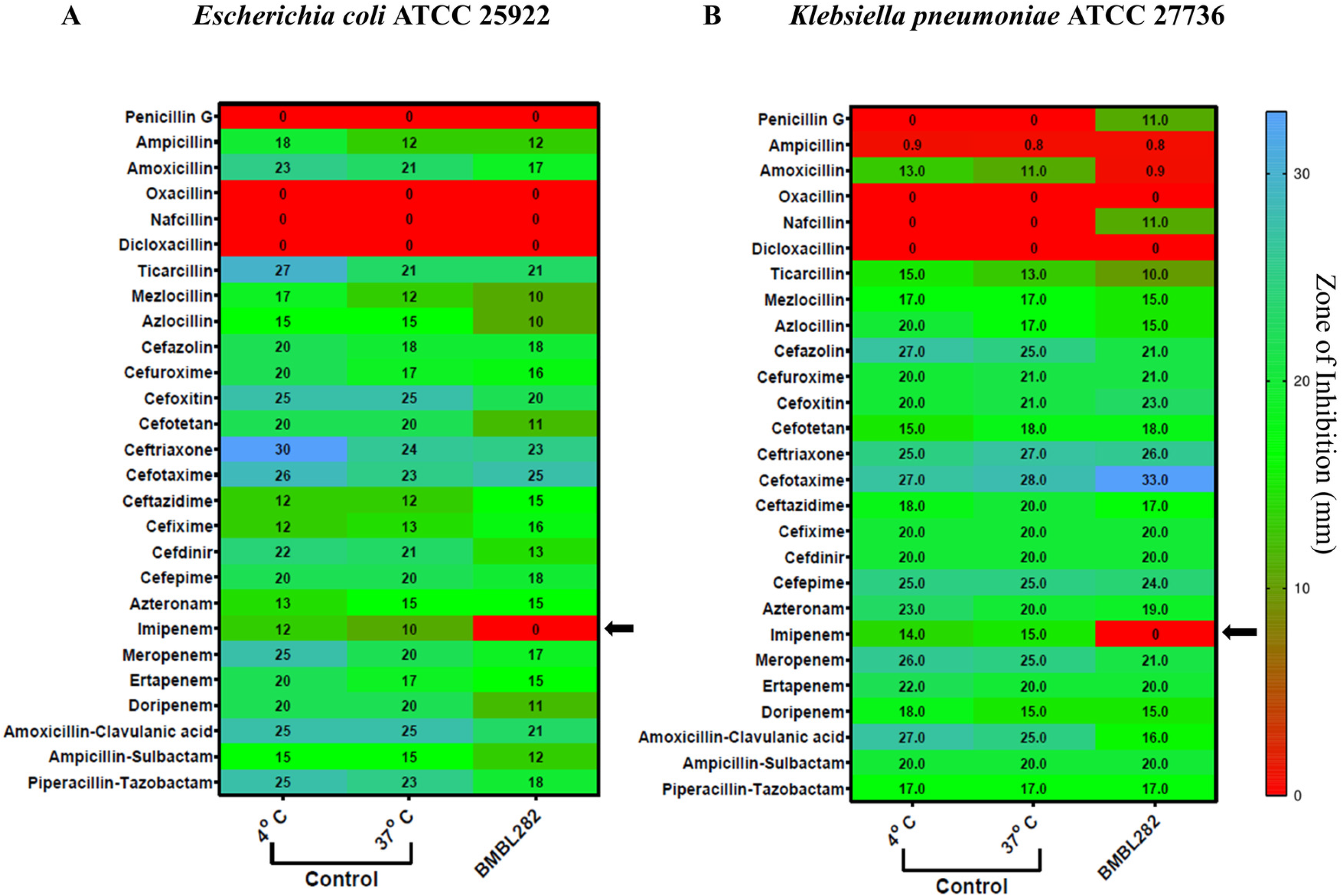
Heat map showing β-Lactam antibiotic degrades ability of BMBL282 (A) Various β-Lactam antibiotics loaded disc (10µg/disc) are incubated with BMBL282 for 12-hours and then placed on lawn of *E. coli* ATCC 25922 (A) or *K. pneumoniae.* (B), followed by measuring the zone of inhibition fresh untreated disc (positive control), disc incubated at 37℃ (experimental control) and with BMBL282 (+P). The best representing experiment from three independent experiment is presented here and the data was interpreted as per CLSI guidelines.

### Agar diffusion assay with Agrobacterium tumefaciens A136 andChromobacterium violaceum 026

BMBL282 *in silico* analysis indicated that it may be a AHL degrading enzyme lactonase. Hence, to investigate the AHL degradation ability of BMBL282, short and long chain AHL were incubated before adding to the *Agrobacterium tumefaciens* A136 and *Chromobacterium violaceum* 026 as biosensor strains (44). These results clearly showed that BMBL282 does not degrade short- and long-chain AHL (Supplementary Fig. S3*A*). This assay is based on the principle that violacein pigment production in *C. violaceum* is inhibited if the AHL is cleaved. Our study shows that AHL molecule upon incubation with BMBL282 did not inhibit violacein production indicating that BMBL did not cleavage the AHL molecules. Most MBL use Zn^2+^ as a cofactor, and *in silico* BMBL282 protein model showed the enzyme possibility of having two Zn^2+^ atoms in HXHXDH domain. Therefore, to rule out this influence, the assay was done in the presence and absence of Zn^2+^. The results showed that BMBL282 could not degrade any type of AHL in the presence and absence of 100 µM ZnCl2.

Furthermore, to eliminate biosensor-based assay limitations and complement the results obtained in the agar diffusion assay, a normalized β-galactosidase activity assay was performed (45, 46). Corroborating previous observations, BMBL282 did not degrade any AHL molecules in the presence or absence of zinc ions. In conclusion, normalized β-galactosidase activity and agar diffusion assay results clearly showed that recombinant BMBL 282 could not degrade AHL molecules (Supplementary Fig. S3B). Hence it is less likely to have any antiquorum-sensing activity (Supplementary Fig. S3A & *B*).

### Determination of fractional inhibitory concentration (FIC) and FIC Index (FICI)

Based upon the structural alignment data, lack of antiquorum sensing activity, the next logical option was to explore BMBL282 metallo beta-lactamase activity against beta-lactam antibiotics. For this, we incubated the beta-lactam class antibiotics Penicillins (Penicillin G, Ampicillin, Amoxicillin, Oxacillin, Nafcilin, Dicloxacillin, Piperacillin, Ticarcillin, Mezlocillin, Azocillin), cephalosporin (Cefazolin, Cephalexin, Cefepime Cefuroxime, Cefoxitin). Cefotetan Ceftriaxone, Cefotaxime, Ceftazidime, Cefixime, Cefdinir), Monobactams (Aztreonam), carbapenems (Imipenem, Meropenem, Ertapenem, Doripenem), and β-lactam inhibitor penicillin combinations (amoxicillin-clavulanic acid, Piperacillin-Tazobactam, Ampicillin-Sulbactam) loaded discs (10 μg/disc) with BMBL282 for 12 h, followed by disk diffusion assay against E*.coli* ATCC 25922 and *Klebsiella pneumoniae* ATCC 27736 bacteria (Fig. 3A & B). Susceptibility testing was performed according to CLSI guidelines (Supplementary Fig. S4). The disc diffusion assay clearly showed that BMBL 282 could only cleave imipenem. The other tested antibiotic did show any significant drop in their antimicrobial activity upon treating with BMBL 282.

To further understand and validate the imipenem-degrading activity, antagonistic activity was performed with BMBL282 in the presence of imipenem or meropenem. Based on the FIC index values, the interaction between the two compounds was determined to be synergistic (FIC<0.5), indifferent (FIC 0.5-4) or antagonistic (FIC> 4). BMBL 282 showed antagonistic effect against Imipenem with FICI values of 16.08 and 8.01 in the presence of *E. coli* and *K. pneumoniae*, respectively. In contrast, BMBL 282 showed an indifferent effect, with FIC values of 2 and 2 in the presence of *E. coli* and *K. pneumoniae*, respectively, against meropenem. With this input, our focus moved to assess the BMBL282 imipenem degrading kinetics. The kinetic characterization of the enzyme demonstrated a maximum velocity (Vmax) of 7.411 × 10⁻⁹ mol s⁻¹ and a Michaelis constant (Km) of 0.50 mM. The turnover number (kcat) was determined to be 2.52 s⁻¹, while the catalytic efficiency (kcat/Km) was calculated as 5.04 × 10³ M⁻¹ s⁻¹.

Most MBL family members are metalloenzymes and exhibit zinc-dependent, typically zinc, i.e., for breaking the β-lactam ring in antibiotic molecules. Interestingly these enzymes are not inhibited by β-lactamase inhibitors but are inactivated in the presence of metal chelators such as ethylenediaminetetraacetic acid (EDTA) (47, 48).Further, we found seven zinc binding site residues of the BMBL282 protein (His 85, His 87, Asp 89, His 90, His174, Asp 193 and His 246) using the Hotspot Wizard tool.

In Gram-negative bacteria, MBL is synthesized and translocated to the membranes through a second system in an unfolded state. The MBL then undergoes final folding and Zn^2+^ acquisition in the periplasm. The role concentration and function of Zn^2+^ metal-binding sites in MBLs have been extensively studied for quite a long time (49). Some studies have shown that MBL is kinetically labile and can acquire Zn(II) ions from the environment or from other sources within 5 min (47).

To further confirm the results, we did direct spectrophotometric measurement of imipenem degradation byBMBL282 in the presence of Zn^2+^, EDTA, and their combination (Fig. 4A). The results clearly showed that even in the presence of Zn^2+^ and EDTA, BMBL 282 could degrade imipenem. To further validate these results, an MIC assay was performed in the presence of 6.8 μg/ml Zn^2+^ and 250 ug/ ml of BMBl 282. As expected, we observed that the MIC of imipenem from <0.78 μg/ml shifted to 12.5 μg/ml, whereas in the presence of EDTA, the MIC value reverted back to <0.78 μg/ml. These data clearly demonstrate that BMBL282 has an imipenem-degrading ability (Fig. 4*A*) whose activity is enhanced in the presence of zinc and addition of the EDTA inhibited the enzyme activity (Fig. 4b).

**Fig 4:**
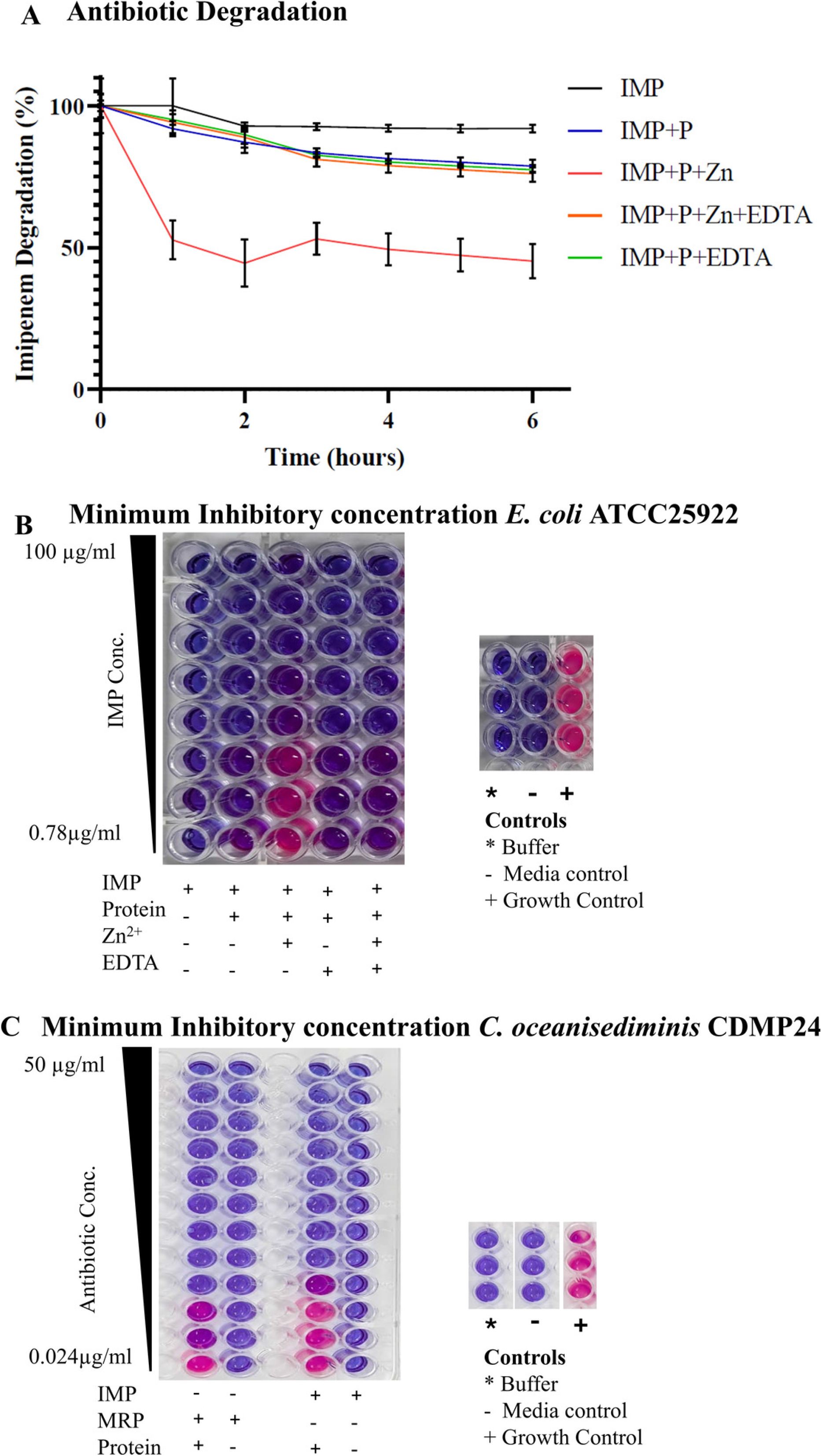
Degradation of imipenem by BMBL282 (A) Shift in the imipenem MIC values was measured in the presence and absence of BMBL282, Zn^2+^ (6.814µg/ml) EDTA (37µg/ml) alone and combinations against *E.coli* and *C.oceanisediminis*. The MIC of imipenem and increased from 0.78 µg/ml (control) to 12.5µg/ml in the presence of Zn^2+^ and BMBL282. (B) Graph showing imipenem degradation in optical density (OD) at 300 nm under different experimental conditions. Data are presented as mean ± SD (N = 3). Statistical analysis was performed using one-way ANOVA followed by Šidák’s multiple comparisons test, with IMP+P+Zn used as the control group. Asterisks (****) indicate statistically significant differences (p < 0.0001) relative to the control.

As a background check, we tested the inhibitory effects of the antibiotics on C. *oceanisediminis*. The results clearly show that C. *oceanisediminis* was inhibited by the tested antibiotics. To confirm the data, we further incubated the imipenem and meropenem disc in BMBL282 before carrying out radial diffusion assay against C. *oceanisediminis*. The results indicated BMBL282 degraded imipenem, hence there is no zone of inhibition, on contrary, there is zone of zone of inhibition for meropenem. To augment further, we did an MIC assay against *C. oceanisediminis* and the results clearly show that the MIC values increase 3- fold higher i.e., 0.024 to 0.195 μg/ml in the presence of BMBL282. All these experiments clearly indicated BMBL282 could be a cryptic gene, as it is not expressed in the *C. oceanisediminis* even in the presence of the antibiotic (Fig. 4c). Given the C. *oceanisediminis* susceptibility to imipenem, it is more likely that the BMBL282 may be a cryptic IMP gene that is not expressed or have other function which needed to be explored (Supplementary Fig. S5).

### Molecular docking studies

The three dimensional (3D) structures of the BMBL282 of *C. oceanisediminis* were not available in the Protein Data Bank (PDB). Hence, it was necessary to model the 3D structure of the proteins for the molecular docking studies. The target proteins BMBL282 of *C. oceanisediminis* sequences were retrieved from the UniProt Database (**A0A160M6H6**). The output of docking results confirmed the molecular interaction between the BMBL282 and imipenem with a binding energy of-5.31 kcal/mol (Fig 5C).

**Fig 5:**
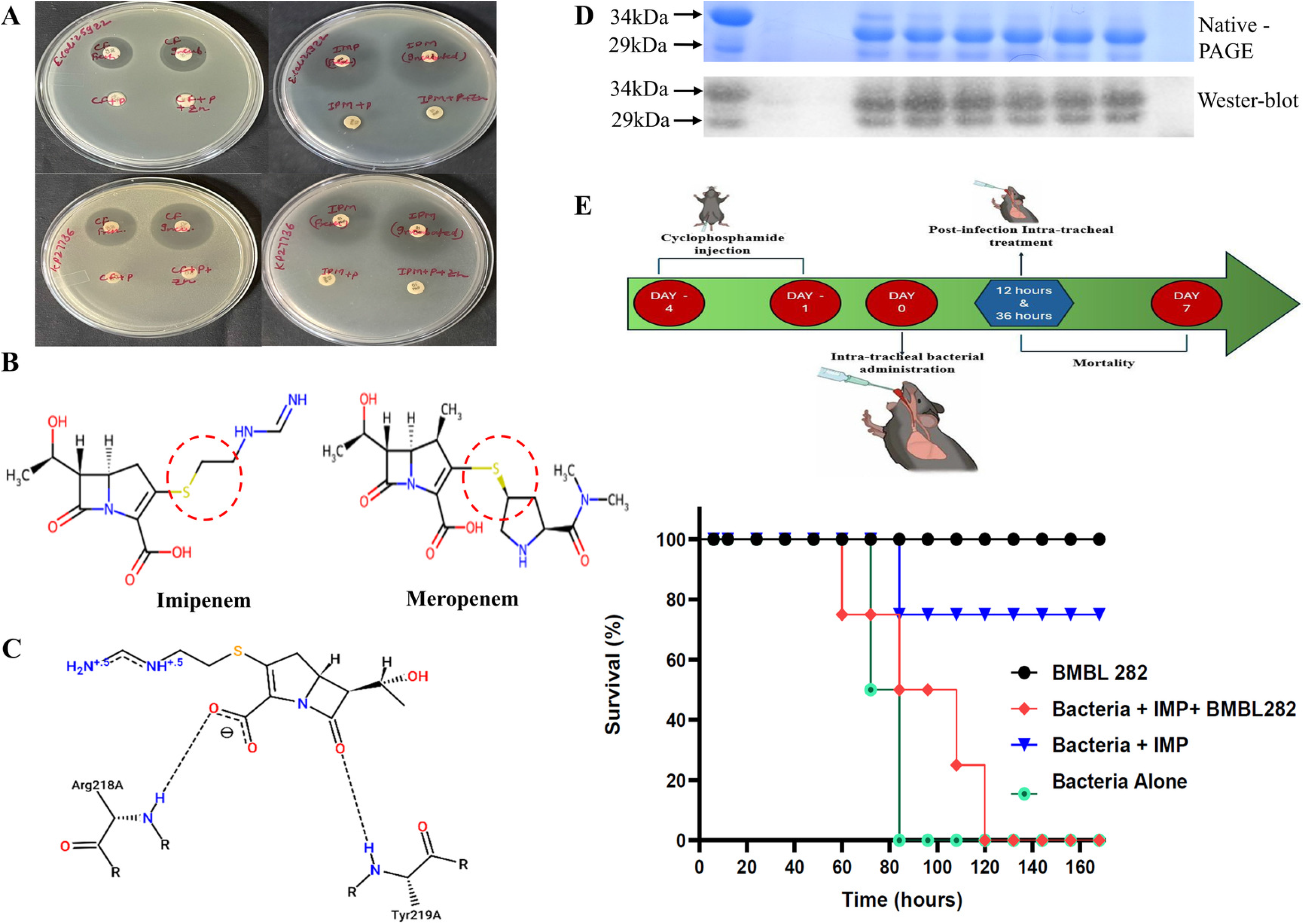
*In-vivo* activity of BMBL282 (A) Image of zone of inhibition showed by indicated disc incubated with or without BMBL282 for 12-hours, followed by placing them on lawn of *E. coli* ATCC 25922 (upper) and *K. pneumoniae* ATCC 27736 (middle) strains and further incubation at 37℃ for 12 hours. (B) the potential cleavage site of imipenem is shown in circle. (C). Docking interaction of BMBL282 with Imipenem. (D) Detection of BMBL282 in the lung lavage using anti-His-tag mAb in western Blot analysis (Lower panel) (E) The survival graph of mice Balb/C infected with *K. pneumoniae* ATCC 27736 in lung, followed by imipenem treatment, in the presence and absence of BMBL282. BMBL282 alone was considered as sham.

### *In vivo* efficacy of BMBL282

It can be argued that BMBL 282 may have imipenemase activity under *in vitro* conditions, which may not translate under *in vivo* conditions. To address this question, we assessed the stability of BMBL282 in lung lavage. The results indicated that BMBL282 is stable upto 5 hours in the lungs (Fig 5D). Given this encouraging results we investigated the *in vivo* activity of imipenem (5 mg/kg) in the presence or absence of BMBL282 (250 µg/kg) in a lung infection survival model. *Klebsiella pneumoniae* ATCC 27736 was injected into the lungs of mice via intratracheal administration. The enzymes BMBL282, Imipenem alone, and imipenem in combination were also administered via the intratracheal route at 12 and 36 h post-infection with appropriate relevant experimental controls, and Kaplan-Meier survival analysis was used to compare the differences between the various groups of results.

As anticipated, all the mice in the control and Imipenem -BMBL282 enzyme treated group died within 72 h (i.e., 3 days) of infection, whereas the group treated with imipenem alone survived until 160 h (i.e., 6 days). In contrast, BMBL282 without infection and sham group mice survived until the end of the experiment (Fig. 5E). We believe that BMBL282 might have cleaved imipenem, thus allowing the bacteria to infect the lungs and leading to mortality. Corroborating the entire data, we conclude that BMBL282 is novel class of MBL with specificity only for imipenem.

## Discussion

Most enzymes from MBL family are promiscuous in nature, with activities against a wide range of substrates, because they contain a stable, highly ancient conserved HXHXH domain (50). Consistent resistance against the “last-resort” antibiotics of the carbapenem family, such as doripenem, imipenem, meropenem, and ertapenem, is often based on the presence of MBL enzymes, as has been repeatedly shown in clinical samples (7, 8, 51). Although the origin of MBL is not well understood and clear, given the menace these enzymes create in the clinic setup, MBLs from terrestrial bacteria have been well-documented and analyzed (43). There is no information on cryptic MBL genes, particularly those from marine bacteria. This study demonstrated the presence of cryptic MBL in the marine bacteria *C. oceanisediminis*.

Some bacteria, especially those from environmental sources, ubiquitously carry the MBL gene either on the chromosome or on plasmids. Both the whole genome sequence deposited at NCBI (Genome ID ASM783023v1) and our studies confirm that *Cytobacillus* bacteria carry the MBL gene on the chromosome (Supplementary Fig. S6). Chromosomal MBL have been reported from the species like *Elizabethkingia meningoseptica*, *Stenotrophomonas maltophilia* (51, 52).

The metallo-beta-hydrolase superfamily consists of a diverse set of hydrolytic enzymes with diverse biological functions, including lactonases and metal beta-lactamases (MBLs). MBL family consist of number of enzymes with different substrate specificity and widely distributed across bacteria, archaea, and eukaryotes. Studies on these enzymes have indicated that the HXHDH domain is essential for catalytic activity and metal coordination, as any mutation in these domains leads to loss of function (53). The HXHXDH domain (where X can be any amino acid except histidine) is ancient and is reported to be present in various enzymes that play a role in diverse biological processes, such as quorum sensing or bacterial communications, RNA processing and DNA repair, suggesting a common ancestral origin for these diverse enzymes (43).

The MBLs family has been divided into four molecular classes, A, B, C, and D, by Ambler, based on the amino acid sequence. Classes A, C, and D carry serine residues at the active site, whereas Class B requires Zn^2+^ as a cofactor (54) and is further classified into subclasses B1 and B2. B1 class MBLs has three histidines and one cysteine, whereas class B2 has asparagine instead of a histidine at the first position of the MBL domain. To accommodate newly emerging MBLs, a separate group called group 3 was proposed based on the substrate profiles, susceptibility to EDTA inhibition, and lack of inhibition by serine beta-lactamase inhibitors. This group is further subdivided into B3a and B3b, based on the broad and narrow substrate specificity, respectively, and poor hydrolysis of carbapenem compared to other substrates (55).

To date, imipenemase homologues from *C. oceanisediminis* have not been structurally or functionally characterized. Structural analysis showed that BMBL 282 is similar to *P. aeruginosa* imipenamase, the phylogenetic analysis of BMBL282 did not cluster with known MBLs and significantly differed from IMP, VIM, and NDM (Supplementary Fig. S6B). To explore potential structural relationships, a PDB-BLAST search was conducted using the amino acid sequence of BMBL282 (NCBI accession: AND37992.1; UniProtKB: A0A160M6H6). The analysis revealed sequence similarity to several members of the metallo-β-lactamase (MBL) superfamily from both marine and terrestrial organisms. Representative homologues with the highest similarity from each subclass (B1–B3) were selected for comparative analysis: metallo-β-lactamase L1 (subclass B3) from *Stenotrophomonas maltophilia* (PDB ID: 7ZO2; 23% identity)(56), metallo-β-lactamase IMP-1 (subclass B1) from *Serratia marcescens* (PDB ID: 6LBL; 21% identity)(57) and carbapenemase ImiS (subclass B2) from *Aeromonas hydrophila* (PDB ID: 3F9O; 17% identity)(58). Multiple sequence alignment of BMBL282 identified a relatively conserved putative Zn²⁺-binding motif (85HXHXDH90). Among the predicted zinc-binding residues, His87, Asp89, and His246 were conserved across the analyzed homologues, Asp193 showed no conservation among the selected reference enzymes. Additional motifs commonly associated with MBL superfamily proteins, including 245GHG247 and 171TPGHTPGH178, were comparatively less conserved (**Fig. 6**). Overall, the sequence alignment indicates that although BMBL282 is evolutionarily divergent, it retains key catalytic residues characteristic of metallo-β-lactamases.

**Fig 6:**
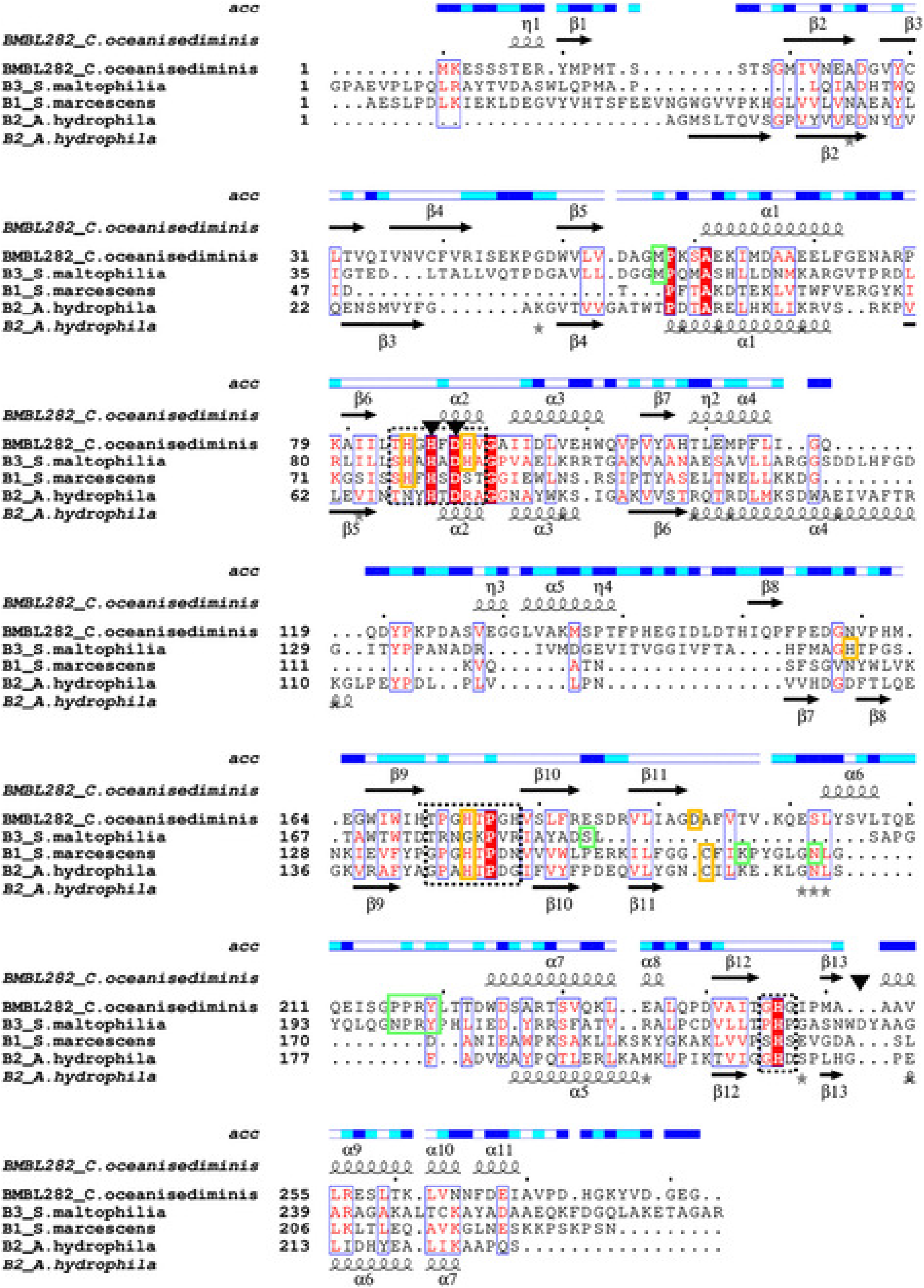
Multiple sequence alignment of BMBL282 with subclass B metallo-β-lactamase homologues. Metallo-β-lactamase L1 (subclass B3) from *Stenotrophomonas maltophilia* (PDB ID: 7ZO2; 23% identity; Hinchliffe *et al.*, 2023), metallo-β-lactamase IMP-1 (subclass B1) from *Serratia marcescens* (PDB ID: 6LBL; 21% identity; Wachino *et al*., 2020) and carbapenemase ImiS (subclass B2) from *Aeromonas hydrophila* (PDB ID: 3F9O; 17% identity; Bebrone *et al.,* 2009). Conserved Zn²⁺-binding residues are marked by black triangles (▼), non-conserved Zn²⁺-binding residues by orange boxes, key active-site residues of respective enzymes by green boxes, and MBL signature motifs by dotted boxes. Secondary-structure elements were derived from the BMBL280 docking model (above) and the carbapenemase ImiS from *A. hydrophila* (PDB ID: 3F9O; below). Residue surface accessibility (acc.) was generated using ESPript 3.0. (https://espript.ibcp.fr).

Studies have shown that, in bacterial species, different bacterial enzymes can evolve separately to hydrolyze the same substrate i.e., antibiotic or drug using a convergent evolution mechanism (49, 59). Kesari et al. showed that MBL has evolved in a convergent manner, not by amino acid substitution, thus leading to variation at sequence levels but not at the functional (59, 60). This observation makes activity assays more important for understanding and confirming protein function in this case. Hence, it was decided to explore the function of BMBL282 using *in vitro* and *in vivo* assays.

Interestingly, BMBML282 has only three histidine residues and no cysteines or asparagine in the domain, showing susceptibility to EDTA, but not to serine beta-lactamase inhibitors i.e antibiotics in combination with clavulanic acid, sulbactam. In the antibiotic disk diffusion assay, C. *oceanisediminis* showed susceptibility to imipenem, but recombinant BMBL282 clearly cleaved imipenem, as demonstrated in various assays (Fig. 3, 4). The MIC assay results confirm the disc assay results, by increasing the MIC value more than 3 fold in comparison to non-treated/absence of BMBL282. This brings us to the next question: Why do bacteria carry cryptic MBL? Currently, two views prevail regarding the evolution of these genes: first, they might encode proteins that perform cellular functions which are yet to be discovered. Second, the constant exposure to antibiotics might have led to the acquisition and maintenance of the genes.

The preservation of metal-binding motifs, together with subclass-specific variations, supports its classification as a Class B metallo-β-lactamase, with closest resemblance to subclass B3 enzymes. However, further structural and functional characterization studies are needed to confirm the class.

To determine the probable mode of action, imipenem and is structurally closet anaglogue meropenem were subjected to cleavage by BMBL282. After the incubation, the antibiotics were tested for their ability to inhibit the *E.coli* or *K. pneumoniae* bacteria. The results show that BMBL282 can cleave the imipenem only, but not meropenem (Fig. 5B). As both meropenem and imipenem have the lactam ring, but differ in the tail region, we are of opinion that BMBL282 might cleave after the sulfur atom attached to the β-lactam ring (Fig. 5C). The inability of BMBL282 to cleave the other beta-lactams antibiotics or AHL molecules, further confirms our hypothesis. Nevertheless, further studies are needed to confirm the cleavage site within the imipnem using the cutting edge technologies.

Interestingly, we observed a domain, “TPGH,” which is frequently found in proteins that play a role in signal transduction, protein-protein interactions, and potential phosphorylation sites (61, 62). However, the exact role of this domain in a particular protein needs to be elucidated, as nothing is known about this domain.

Pleiotropic proteins with MBL folds have been shown to exhibit promiscuous enzymatic activities. Gene and protein sequence analysis using *in silico* tools indicated that the BMBL 282 protein belongs to the beta hydrolase family. Most enzymes from the metallo-hydrolase family are promiscuous in nature, with activities against a wide range of substrates because they contain a stable, highly ancient, conserved fold. Hence, it is possible that BMBL282 may cleave other substrates also. The limitation of the present study is that we did not explore other potential beta-hydrolyase substrates, such as RNA. We believe that analyzing these in future studies could broaden our understanding of BMBL282.

Given the present study, it is tempting to speculate that cryptic genes present in the marine genome may be a source of new antibiotic resistance genes. Future studies will aim to explore cryptic genes with HXHXDH domains in other marine bacteria to determine whether they show narrow- or broad-spectrum antibiotic degradation. We believe that a critical understanding of the cryptic antibiotic resistance genes is important in our fight against the global silent pandemic of antibiotic resistance.

Using *in silico*, *in vitro* and *in vivo* assay results, we demonstrated that the cryptic gene BMBL282 in *C. oceanisediminis* encodes a enzyme that can cleave imipenem, does not share homology with the other reported IMP genes but has two characteristic domains: HXHXDH and TPGH. To the best of our knowledge, this is the first report of presence of cryptic gene in the Marine bacterium *C. oceanisediminis* that can degrade imipenem but not the other antibiotics of the carbapenem family.

## Acknowledgments

MP is grateful to the Ex-Director(S), CDRI, Lucknow, India, CSIR New Delhi [IHP0019, IHP0016, MLP0107], DST [CRG/2022/005456], ICMR [Em/Dev/SG/57/1654/2023-e-office-176936], and Ignite Life Science Foundation [ACORN-AMR(2)/2024/003] for providing financial assistance. PG is grateful to CSIR (31/004(1365)/2019-EMR-I), New Delhi, India. The authors thank Mr. Angney Lal and Mr. Atul Krishna for their technical assistance during the study. CSIR-CDRI communication number

## Supplementary figures

Fig S1: - *Cytobacillus oceanisediminis* (CDMP24) isolation. (A) Photograph representing the marine water sampling site Gulf of Mannar, India. (8°28′N 79°01′E / 8.47°N 79.02°E). (B) Bacterial pellet of *Cytobacillus oceanisediminis.* (C) Gel image showing DNA isolation of *Cytobacillus oceanisediminis* (CDMP24) (Left), (Right) lane1: - Marker, Lane2&3 PCR amplified product. (D) DNA sequence of *Cytobacillus oceanisediminis* (CDMP24) strain 16S rRNA partial sequence (submitted to KU933344.1).

Fig S2: - Cytobacillus oceanisediminis Lactonase282 (BMBL282)- (A) Gene sequence (B) SDS PAGE analysis of expressed protein (BMBL282) from *Cytobacillus oceanisediminis*. (B) The gel pic showing the purification of BMBL282 (34kDA) protein by Ni-NTA column. Lane 1-Whole cell lysate, Lane 2- Flow through, Lane 3- Wash buffer, Lane 4- desired purified protein (BMBL282), Lane 5- Purified and dialyzed protein and Lane 6- Protein marker. (C) Western blot of Ni-NTA affinity purified protein. Bands observed via anti-His antibody labeling. Lanes 1 represents protein marker and Lane 2 represent eluted fraction from Ni NTA column respectively.

Fig S3: - Anti-quorum sensing activity of BMBL282. (A) Agar diffusion assay with Chromobacterium violacium 026 (left plate): (+) control: Absence of C6 HSL molecule, (-) negative control: Non-degraded AHL Molecule. Agar diffusion assay with Agrobacterium tumefaciens A136 (right plate): (+) control: Absence of C14-HSL, (-) negative control: Non-degraded AHL Molecule. Lower left column show the results obtained Anti-quorum sensing activity of BMBL282 when different AHL molecules were used. (B) Normalised β-galactosidase activity with Agrobacterium tumefaciens A136 against different AHL molecules incubated with or with Protein (BMBL282) (C) Whole drop images of BMBL-282 with micro-crystals.

Fig S4: - *In-vitro* Radial disk diffusion assay of *Escherichia coli* ATCC 25922 (left) and *Klebsiella pneumoniae* KP27736 (right) using β-lactam antibiotics. Three conditions were tested: (1) antibiotic disc alone, (2) antibiotic disc incubated without enzyme (control), and (3) antibiotic disc incubated with purified protein BMBL282 for 12-hours, followed by placing them on lawn of *E. coli* ATCC 25922 bacteria and further incubation at 37℃ for 12 hours. Zones of inhibition were measured to assess the potential β-lactamase activity of BMBL282.

Fig S5: - Results of Radial disk diffusion assay of *Cytobacillus oceanisediminis* using β- lactam antibiotics disc incubated with purified protein BMBL282. (A) table and figure showing results of zones of inhibition in the absence of the BMBL282. (B) Figure showing results of zones of inhibition with the imipenem and meropenem antibiotic disc alone, treated and untreated (upper panel) blownup images (lower panel) of the imipenem (IMP) and meropenem (MRP) antibiotic disc, and imipenem and meropenem antibiotic disc incubated with purified protein BMBL282 for 12-hours, followed by placing them on lawn of *Cytobacillus oceanisediminis* bacteria and further incubation at 37℃ for 12 hours.

Fig S6: - MIC assay of imipenem against *E. coli* ATCC 25922 in the presence and absence of recombinant protein BMBL282. Assays were performed with ZnCl₂ (6.814 µg/mL) or EDTA (37 µg/mL) supplementation. The MIC of imipenem increased from 0.78 µg/mL (control) to 12.5 µg/mL in the presence of Zn²⁺ and BMBL282, indicating zinc-dependent hydrolysis consistent with metallo-β-lactamase activity.

Fig S7: - MIC assay of imipenem and meropenem against Cytobacillus oceanisediminis CDMP 24 in the presence and absence of recombinant protein BMBL282.Minimum inhibitory concentrations (MICs) were determined to assess the effect of recombinant protein BMBL282 on carbapenem susceptibility. In the absence of protein, both imipenem (IPM) and meropenem (MRP) showed MIC values of <0.024 µg/mL. In the presence of recombinant BMBL282, the MIC of imipenem increased to 0.195 µg/mL and the MIC of meropenem increased to 0.048 µg/mL, indicating reduced susceptibility of C. oceanisediminis CDMP 24 to carbapenems in the presence of the protein.

Fig S8: - Degradation of imipenem by BMBL282 over a period of 6 hours.

Fig S9: Complete gels of SDS–PAGE and Western blot analysis of BMBL282 stability under protease and lung lavage treatment. Lane 1,Protein Ladder; Lane 2, untreated BMBL282 (34kDa, control); Lane 3, trypsin-treated BMBL282; Lane 4, proteinase K-treated BMBL282; Lanes 5–10, BMBL282 incubated with mouse lung lavage fluid for 1, 2, 3, 4, 5, and 6 h, respectively; Lane 11, lung lavage fluid alone (negative control); Lane 12, His-tag positive control.

Fig S10: - Confirmation of location of BMBL282 cryptic gene (A) The agarose gel electrophoresis image shows the amplification and analysis of the BMBL282 gene from *CDMP24*. Lane 1 contains the DNA ladder, serving as a molecular weight marker. Lane 2 displays the genomic DNA extracted from *CDMP24*, while lane 3 shows the plasmid DNA isolated from the same strain. Lane 4 represents the PCR negative control, which was set up without any template DNA to confirm the absence of contamination. Lane 5 shows the PCR amplification product of the BMBL282 gene amplified from the genomic DNA of *CDMP24* using specific primers BMBL282F and BMBL282R. Lane 6 displays the PCR amplification product of the BMBL282 gene from the plasmid DNA of *CDMP24* using the same primer set. (B) Phylogenetic tree analysis of BMBL282 gene. The BMBL282 clustered with MBL’s which were reported to be on chromosome.

**Supplementary Table 1: - Primers Used in this Study.**

